# Inducible Lung Epithelial Resistance Requires Multisource Reactive Oxygen Species Generation to Protect against Bacterial Infections

**DOI:** 10.1101/471649

**Authors:** Hayden H. Ware, Vikram V. Kulkarni, Yongxing Wang, Miguel Leiva Juarez, Carson T. Kirkpatrick, Jezreel Pantaleón García, Shradha Wali, William K. A. Sikkema, James M. Tour, Scott E. Evans

## Abstract

Pneumonia remains a global health threat, in part due to expanding categories of susceptible individuals and increasing prevalence of antibiotic resistant pathogens. However, therapeutic stimulation of the lungs’ mucosal defenses by inhaled exposure to a synergistic combination of Toll-like receptor (TLR) agonists known as Pam2-ODN promotes mouse survival of pneumonia caused by a wide array of pathogens. This inducible resistance to pneumonia relies on intact lung epithelial TLR signaling, and inducible protection against viral pathogens has recently been shown to require increased production of epithelial reactive oxygen species (ROS) from multiple epithelial ROS generators. To determine whether similar mechanisms contribute to inducible antibacterial responses, the current work investigates the role of ROS in therapeutically-stimulated protection against *Pseudomonas aerugnosa* challenges. Inhaled Pam2-ODN treatment one day before infection prevented hemorrhagic lung cytotoxicity and mouse death in a manner that correlated with reduction in bacterial burden. The bacterial killing effect of Pam2-ODN was recapitulated in isolated mouse and human lung epithelial cells, and the protection correlated with inducible epithelial generation of ROS. Scavenging or targeted blockade of ROS production from either dual oxidase or mitochondrial sources resulted in near complete loss of Pam2-ODN-induced bacterial killing, whereas deficiency of induced antimicrobial peptides had little effect. These findings support a central role for multisource epithelial ROS in inducible resistance against bacterial pathogens and provide mechanistic insights into means to protect vulnerable patients against lethal infections.

## INTRODUCTION

Lower respiratory tract infections remain the leading cause of premature death and disability among both otherwise healthy and immunosuppressed people worldwide (1-5). In an era of increasing antimicrobial resistance, human global hypermobility, proliferation of emerging and weaponized pathogens, aging populations, and ever-expanding categories of immunocompromised patients, the acute complications of pneumonia exact a staggering toll, killing an estimated 2.7 million people per year (6-10). The 1943 introduction of penicillin for pneumonia management was a medical triumph (11), but the intervening decades have witnessed escalating age-adjusted pneumonia hospitalization rates (12-14) without survival rate improvements of corresponding magnitude (15). In an effort to address the persisting threat of pneumonia to vulnerable populations, our laboratory has developed a program focused on manipulating the intrinsic antimicrobial capacity of the host to prevent pneumonia.

Based on this program, we have reported that the mucosal defenses of the lungs can be stimulated to protect mice against a wide array of otherwise lethal pneumonias, including those caused by antibiotic-resistant bacteria (16-19). This *inducible resistance* is achieved following a single inhaled treatment comprised of a synergistic combination of Toll-like receptor (TLR) agonists: a diacylated lipopeptide ligand for TLR2/6, Pam2CSK4, and a class C unmethylated 2=-deoxyribocytidine-phosphate-guanosine (CpG) ligand for TLR9, ODN M362 (hereafter, Pam2-ODN) (16, 17, 20).

Inducible resistance against pneumonia requires intact lung epithelial TLR signaling mechanisms, whereas no individual leukocyte populations have been identified as essential to Pam2-ODN-enhanced pneumonia survival (16, 21). Given the epithelial requirement for inducible resistance in vivo (16, 22), we sought to determine whether epithelial cells were sufficient to act as autonomous antibacterial effector cells of therapeutically inducible protection. We recently reported that Pam2-ODN-induced antiviral protection requires therapeutic induction of reactive oxygen species (ROS) via a novel multisource mechanism (23), but it is unknown whether similar processes are required for inducible antibacterial defense.

We report here that Pam2-ODN induces active antibacterial responses from intact lungs and isolated lung epithelial cells that reduce pathogen burden, attenuate infectivity, and enhance survival. Moreover, we find that the protection requires epithelial generation of ROS via dual mechanisms, providing meaningful insights into the mechanisms underlying the novel synergistic interactions observed between the TLR ligands.

## RESULTS

### Pam2-ODN treatment reduces pathogen burden and inflammatory injury in bacterial pneumonia

We have previously reported that a single nebulized treatment with Pam2-ODN results in improved survival of otherwise lethal pneumonias, including those caused by *P. aeruginosa* (16, 17, 20, 21). Here, we found that the protection afforded by Pam2-ODN (Figure 1A) is associated with reduced bacterial burden immediately after infection, whether culturing lung homogenates or bronchoalveolar lavage (BAL) fluid (Figure 1B), suggesting that a Pam2-ODN-induced antimicrobial environment existed at the time of infection. No significant differences were noted in the performance of the two culture methods, in terms of precision or magnitude of induced bacterial reductions by Pam2-ODN, though the absolute bacteria per ml tended to be higher in the BAL-obtained samples than in the lung homogenates. Proportionally similar inducible reductions in pathogen burden were observed both immediately after infection and 24 h after infection when mice were infected with fluorescent *P. aeruginosa*, then their lungs were examined by fluorescence microscopy (Figure S1).

**Figure 1.**
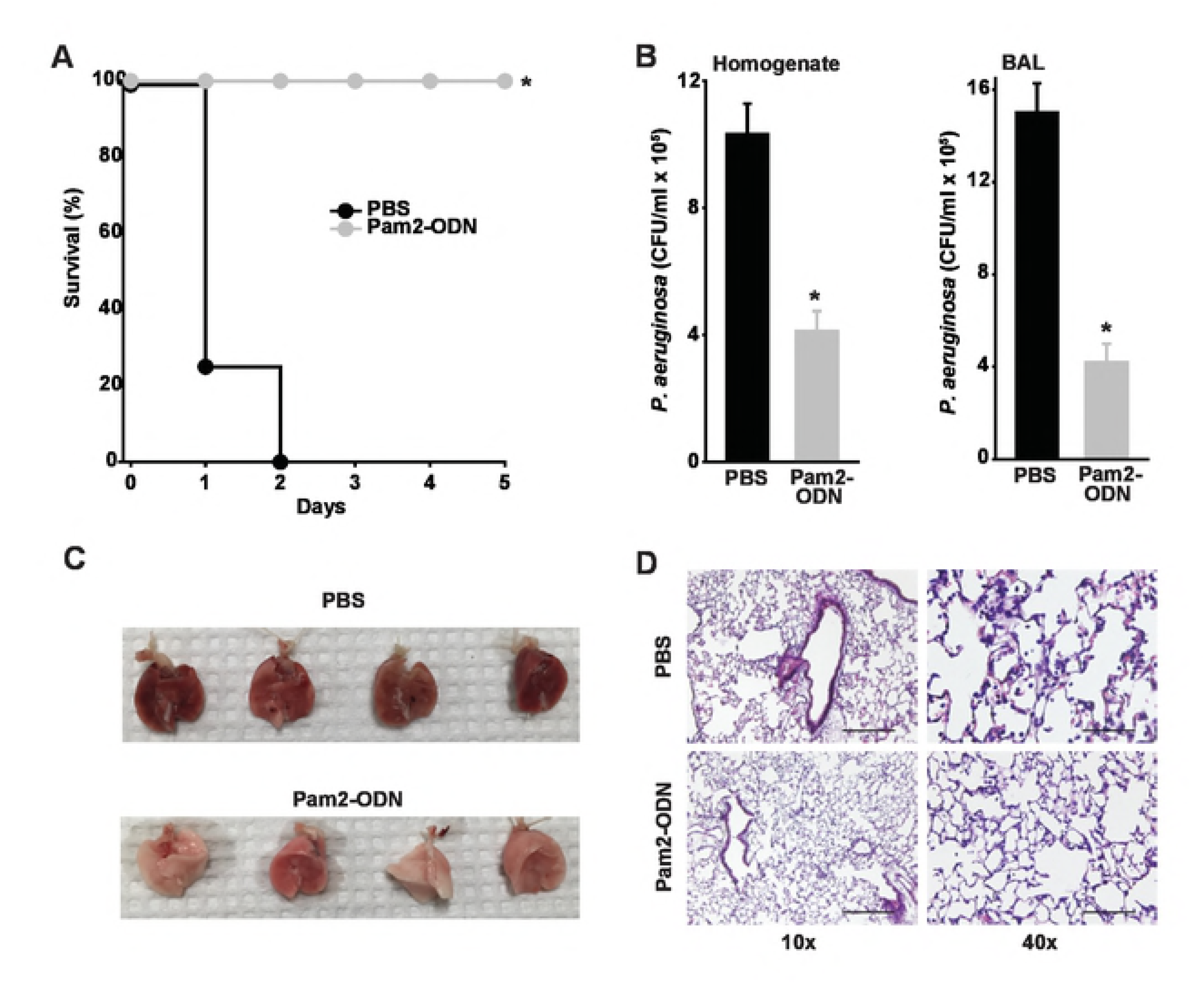
Pam2-ODN protects against bacterial pneumonia. (**A**) Survival of wildtype mice treated with Pam2-ODN or PBS (sham) by aerosol 24 h before challenge with *P. aeruginosa*. (**B**) Pathogen burden of mice in **A** immediately after challenge, as assessed by serial dilution culture of lung homogenates (*left* panel) or BAL fluid (*right* panel). (**C**) Gross appearance of mouse lungs 24 h after *P. aeruginosa* challenge following treatment with Pam2-ODN or sham. (**D**) Hematoxylin-eosin stained histology of lungs in **C**. Scale bar = 400 μm *left* panels, 100 μm *right* panels. Each panels is representative of at least three independent experiments. N = 8 mice/group for survival, N = 4 mice/group for pathogen burden. * p < 0.0002 vs. PBS-treated; ** p < 0.002 vs PBS-treated.

Although inhaled treatment with Pam2-ODN induces transient lung neutrophilia (16), we found here that the antimicrobial environment associated with Pam2-ODN-induced resistance also protected against inflammatory lung injury. Lungs harvested 24 h after *P. aeruginosa* challenge demonstrate severe hemorrhagic pneumonia in sham-treated mice, but there is almost no evidence of such injury in Pam2-ODN-treated mice (Figure 1C). Similarly, histologic inspection of Pam2-ODN-treated lungs 24 h after infection demonstrate substantially less inflammatory cell infiltration and notably fewer bacteria. This is congruent with earlier studies (16) suggesting that the difference in *P. aeruginosa* continues to increase between active and sham treated groups as time elapses, indicating that the antimicrobial environment persists beyond the period of the initial challenge.

### Pam2-ODN induces bacterial killing by isolated lung epithelial cells across a broad concentration and temporal range

Congruent with the *in vivo* observations, we have found that treatment of isolated human or mouse lung epithelial cells results in significant reductions in culture bacterial burdens (16, 17, 21, 22). Based on empiric *in vivo* efficacy optimization, Pam2-ODN is administered in a fixed 4:1 molar ratio (18, 20). Figure 2A-B shows that, when delivered in this ratio, the antibacterial effect is inducible across treatment concentrations that extend to at least a 2 log10 range (Pam2 0.124-12.4 μM; ODN 0.031-3.1 μM). Higher Pam2-ODN treatment concentrations are expected to induce even greater bacterial killing than that shown, but when calculating the estimated deposition of the ligands in 20 μl mouse airway lining fluid (24) or in 10-30 ml of human airway lining fluid (25) after nebulization, it is unlikely that such high concentrations can be achieved in vivo. To avoid presenting responses that are easily detectable but not physiologically relevant to the *in vivo* model, all subsequent figures include data achieved with a lower Pam2-ODN dose (2.23 uM Pam2 and 0.56 uM ODN) that we calculate to be achievable by nebulization, except when labelled as dose response plots.

In this model, bacteria are inoculated into the epithelial cultures in log phase growth and there are no antibacterial leukocyte contributions. So, the antimicrobial epithelial responses must be active very early in the course of infection. To determine how quickly these responses could be induced, we tested treatment intervals prior to challenge and found that the most profound antibacterial responses seemed to be achieved with six or more hours of treatment, but significant bacterial burden reductions were achieved in a much shorter period (Figure 2C-D) in both mouse and human cells. In fact, the antibacterial effect was even observed when Pam2-ODN is administered simultaneous to the infectious challenge.

**Figure 2.**
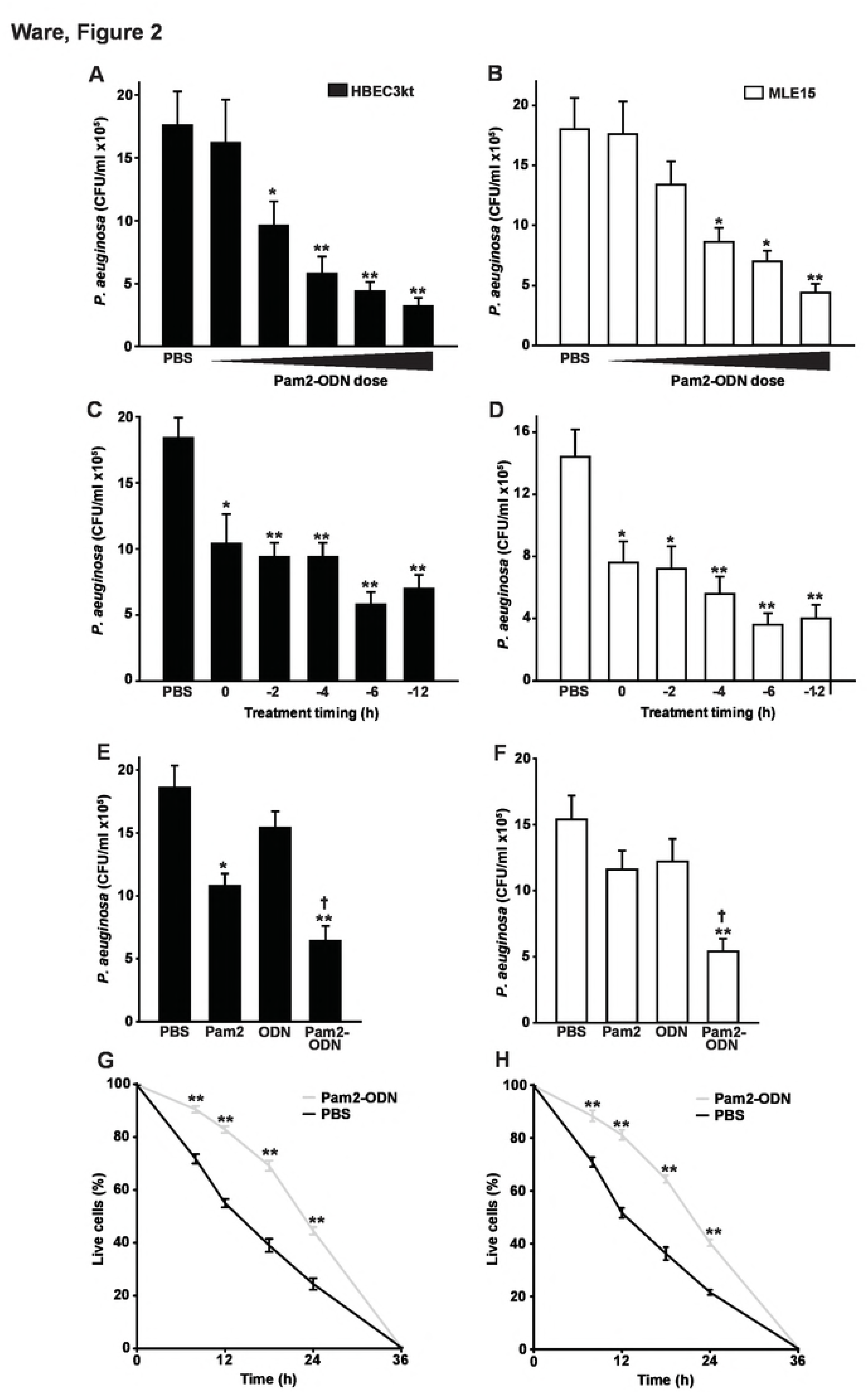
Pam2-ODN induces antibacterial responses in isolated lung epithelial cells. HBEC3kt (**A**) or MLE15 (**B**) cells were treated for 6 h with PBS or escalating doses of Pam2-ODN (range: Pam2 0.12-12.4 uM, ODN 0.03-3.10 uM), then challenged with *P. aeruginosa*. Shown are culture bacterial burdens 6 h after challenge. HBEC3kt (**C**) or MLE15 (**D**) cells were treated for 6 h with PBS or Pam2-ODN (middle dose used in **A** and **B**, 2.23 uM Pam2 and 0.56 uM ODN) for the indicated interval prior to challenge with *P. aeruginosa*. Shown are culture bacterial burdens 6 h after challenge. HBEC3kt (**E**) or MLE15 (**F**) cells were treated for 6 h with the indicated treatments, then challenged with *P. aeruginosa*. Shown are culture bacterial burdens 6 h after challenge. HBEC3kt (**G**) or MLE15 (**H**) cells were treated with PBS or Pam2-ODN for 6 h prior to *P. aeruginosa* challenge. Cell survival determined by Trypan blue exclusion is shown at the indicated time points. Each panels is representative of at least three independent experiments. * p < 0.05 vs. PBS-treated; ** p < 0.005 vs. PBS-treated; † p < 0.05 vs. either single ligand treatment.

### Pam2-ODN interact synergistically to induce bacterial killing

Further substantiating the *in vitro* model as relevant to study of the *in vivo* pneumonia protection associated with Pam2-ODN treatment, we found that the antibacterial effect of the combined Pam2-ODN treatment was supra-additive to the effects of equimolar ligands delivered individually. Pam2 alone induced a modest reduction in bacterial burdens in human epithelial cells (Figure 2E). The magnitude of this effect is similar to the degree of protection we have observed *in vivo* with Pam2 alone (17). ODN alone did not induce any significant reductions in bacterial burden in either human or mouse epithelial cultures (Figure 2F). However, in both models, the combination of Pam2 and ODN resulted in greater anti-pseudamonal effects that the combined effects of the two ligands delivered alone.

### Pam2-ODN extends epithelial survival of *Pseudomonas* infections

While Pam2-ODN induces a robust antibacterial effect, it has not been previously established whether the antimicrobial response was associated with a fitness cost to the cells themselves. For instance, it is conceivable that the microbicidal response might also be toxic to the host cells or it is possible that programmed cell death pathways contribute to bacteriostatic effects. Indeed, we have previously reported that inducible epithelial resistance is correlated with transient but profound induction of inflammatory mediators (16, 17, 20), and here find significant induction of genes for both proinflammatory cytokines and antimicrobial peptides from lung epithelial cells treated with Pam2-ODN (Figure S1A). However, we have not found reduced survival of lung epithelial cells following Pam2-ODN treatment in the absence of infection and see dramatically improved cell survival of viral infections when the cells received Pam2-ODN pretreatment (23). To investigate the effect of Pam2-ODN treatment on epithelial cell survival of bacterial infections, Trypan blue exclusion was used to determine cell viability following *P. aeruginosa* infection. While all epithelial cells were dead by 36 h after the infectious challenge, regardless of pretreatment, both mouse and human epithelial cells lived longer on average and had a greater percentage of cells alive at every intermediate time point if pretreated with Pam2-ODN (Figure 2G-H). These findings support that the antibacterial effect of the epithelial response to Pam2-ODN is more beneficial than any potential fitness cost.

### Pam2-ODN-induced antibacterial effects require DUOX-dependent ROS production

Antimicrobial peptides are well established contributors to lung epithelium-mediated antibacterial defense (22), and genes encoding antimicrobial peptides such as lipocalin 2 and acute phase serum albumin A proteins are some of the most strongly upregulated transcripts following Pam2-ODN treatments of lungs or isolated lung epithelial cells (16, 17). However, *P. aeruginosa* challenge of mice deficient in these antibacterial molecules revealed little defect in Pam2-ODN-inducible protection, even when more than one antimicrobial peptide gene was knocked out Figure S2 B-E). These data suggest that, although they are robustly induced, these individual antimicrobial peptides are not essential effectors of inducible resistance, and prompted investigations into alternate effector mechanisms.

ROS are increasingly recognized to function as direct antimicrobial effector molecules, most likely through lipid peroxidation of microbial membranes and DNA damage, in addition to their well-established roles as signaling molecules (26). We previously hypothesized that ROS contribute to Pam2-ODN-induced epithelial antibacterial effects (16, 18, 19). More recently, we confirmed that ROS are essential to Pam2-ODN-induced antiviral responses and have published a comprehensive characterization of the epithelial ROS species induced by Pam2-ODN treatment (23). Figures 3A-B confirm that Pam2-ODN induces dose-dependent production of ROS from both human and mouse lung epithelial cells, as measured by fluorescence signal from cell permeant carboxy-H2DCFDA. Our previous findings indicate that only hydrogen peroxide and superoxide are demonstrably increased by epithelial treatment (23), and there is no reason to suspect that induction of alternate species is reflected by the carboxy-H_2_DCFDA signal in the current studies using identical culture and treatment models.

**Figure 3.**
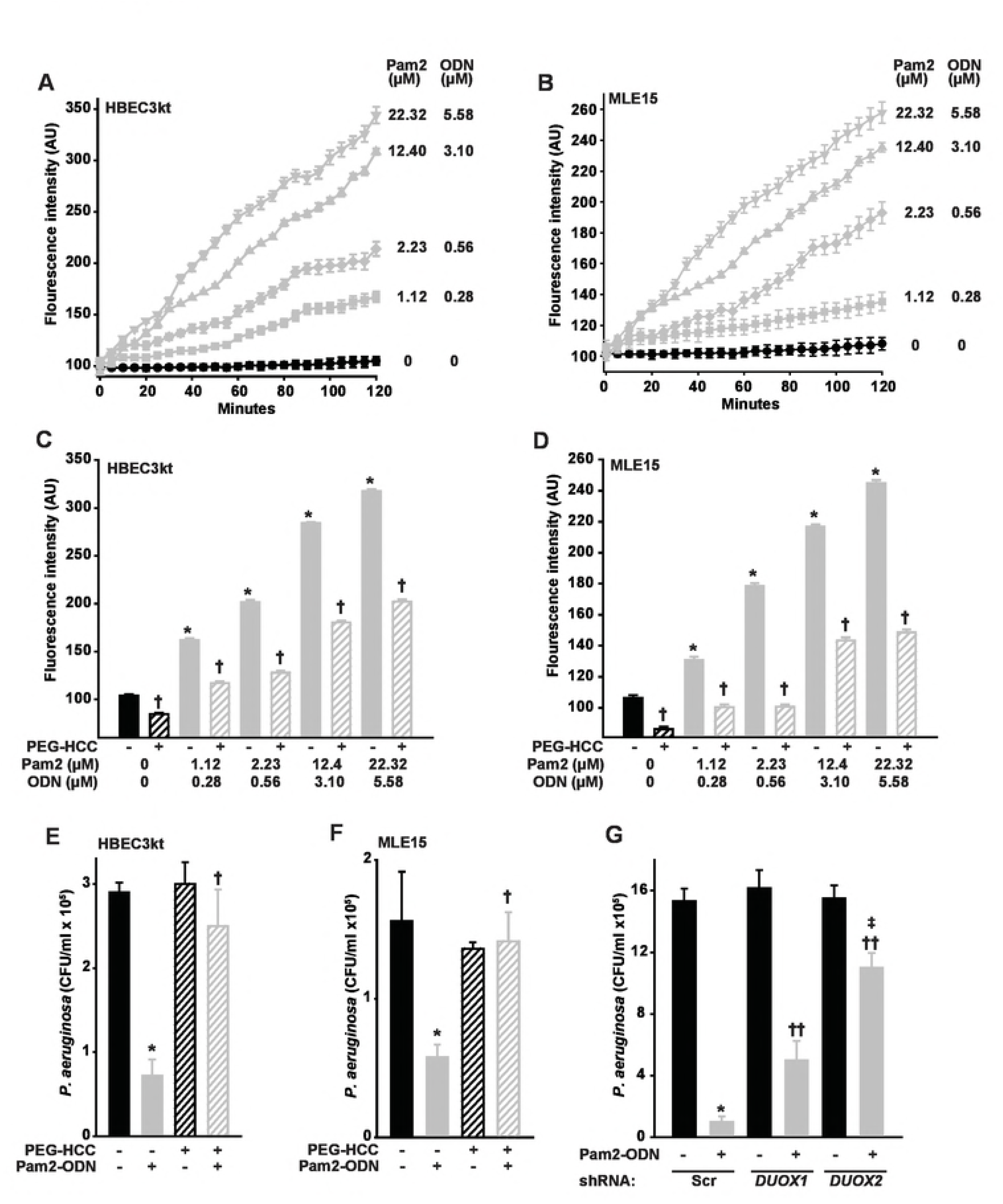
Pam2-ODN induces antibacterial ROS from isolated lung epithelial cells. HBEC3kt (**A**) or MLE15 (**B**) cells were exposed to CO-H_2_DCFDA, treated with the indicated doses of Pam2-ODN, then fluorescence intensity was measured every 5 min. HBEC3kt (**C**) or MLE15 (**D**) cells were pretreated with PEG-HCC or PBS, exposed to CO-H_2_DCFDA, then treated with the indicated dose of Pam2-ODN. Shown are fluorescence intensity 100 min after treatment. HBEC3kt (**E**) or MLE15 (**F**) cells were pretreated with PEG-HCC or PBS, treated for 6 h with PBS or Pam2-ODN (Pam2 2.23 uM, ODN 0.56 uM), then challenged with *P. aeruginosa*. Shown are culture bacterial burdens 6 h after challenge. (**G**) HBEC3kt cells were stably transfected with scrambled (control) shRNA or shRNA targeting *DUOX1* or *DUOX2*, then treated with PBS or Pam2-ODN for 6 h prior to *P. aeruginosa* challenge. Shown are culture bacterial burdens 6 h after challenge. Each panels is representative of at least three independent experiments. * p < 0.005 vs no Pam2-ODN treatment; † p < 0.005 vs no PEG-HCC, same Pam2-ODN; †† p < 0.02 vs scrambled shRNA + Pan2-ODN; ‡ p < 0.003 vs *DUOX1* knockdown + Pam2-ODN.

Acting by superoxide dismutation and radical annihilation (27, 28), application of poly(ethylene glycolated) hydrophilic carbon clusters (PEG-HCCs) (28, 29) to the culture media significantly reduced epithelial ROS, as demonstrated by CO-H_2_DCFDA fluorescence, at all Pam2-ODN doses (Figure 3C-D). Notably, by reducing epithelial ROS, PEG-HCC treatment also significantly impaired the Pam2-ODN-induced epithelial antibacterial effect (Figure 3E-F), supporting a ROS requirement for inducing the protective response from both human and mouse lung epithelial cells.

While all NADPH oxidase (NOX) isoforms are reported to be expressed by lung epithelia, the primary source of lung epithelial ROS are the dual oxidases DUOX1 and DUOX2 (sometimes called NOX6 and NOX7) (30-32). To test the specific requirement for DUOX-derived ROS in Pam2-ODN-induced antibacterial defense, we used shRNA to stably knockdown *DUOX1* and *DUOX2* in HBEC3kt cells, then assessed the effect on Pam2-ODN-induced reductions in influenza burden. Figure 3G shows that knocking down *DUOX1* moderately impairs the Pam2-ODN-induced epithelial antimicrobial response and knocking down DUOX2 severely impairs the inducible antibacterial effect. This is congruent with prior reports that DUOX1 produces a relatively consistent amount of ROS, though this production can be moderately enhanced by IL-4 and IL-13 exposure(33), whereas DUOX2-dependent ROS production can be profoundly increased by activating existing DUOX2 and increasing *DUOX2* and *DUOXA2* transcription following exposure to cytokines such as IFNγ(33). Interestingly, while the *DUOX1* requirement for inducible antipseudomonal defense appears to be less substantial than the DUOX2 requirement, the inducible protection defect observed in DUOX1 knockdown cells is more profound than that observed in virus challenged DUOX1 knockdown cells (23).

### Pam2-ODN-induced antibacterial effects require mitochondrial ROS production

Although we confirmed that DUOX-dependent ROS production is required for the inducible bacterial killing, there is accumulating evidence that mitochondria-derived ROS can also participate in antimicrobial responses of nonphagocytes (23, 34-37). To test whether mitochondrial ROS also contribute to the inducible antibacterial effect of Pam2-ODN, mitochondrial ROS were selectively scavenged with mitoTEMPO prior to *P. aeruginosa* challenge with or without Pam2-ODN pretreatment. Figure 4 A-B shows that mitochondrial ROS scavenging profoundly impaired the Pam2-ODN induced bacterial killing by mouse and human epithelial cells. To address potential off-target effects or nonspecific ROS scavenging of mitoTEMPO, we tested whether we could reduce inducible mitochondrial ROS production, rather than scavenging produced ROS. Figure 4C shows that combination treatment with a mitochondrial complex II inhibitor and a respiratory chain uncoupler reduces mitochondrial ROS production at every tested dose of Pam2-ODN. This impaired Pam2-ODN-induced mitochondrial ROS production resulted in bacterial killing defects that mirrored the mitochondrial ROS scavenging experiments (Figure 4 D-E), revealing a requirement for mitochondrial ROS in Pam2-ODN-induced antibacterial responses.

**Figure 4.**
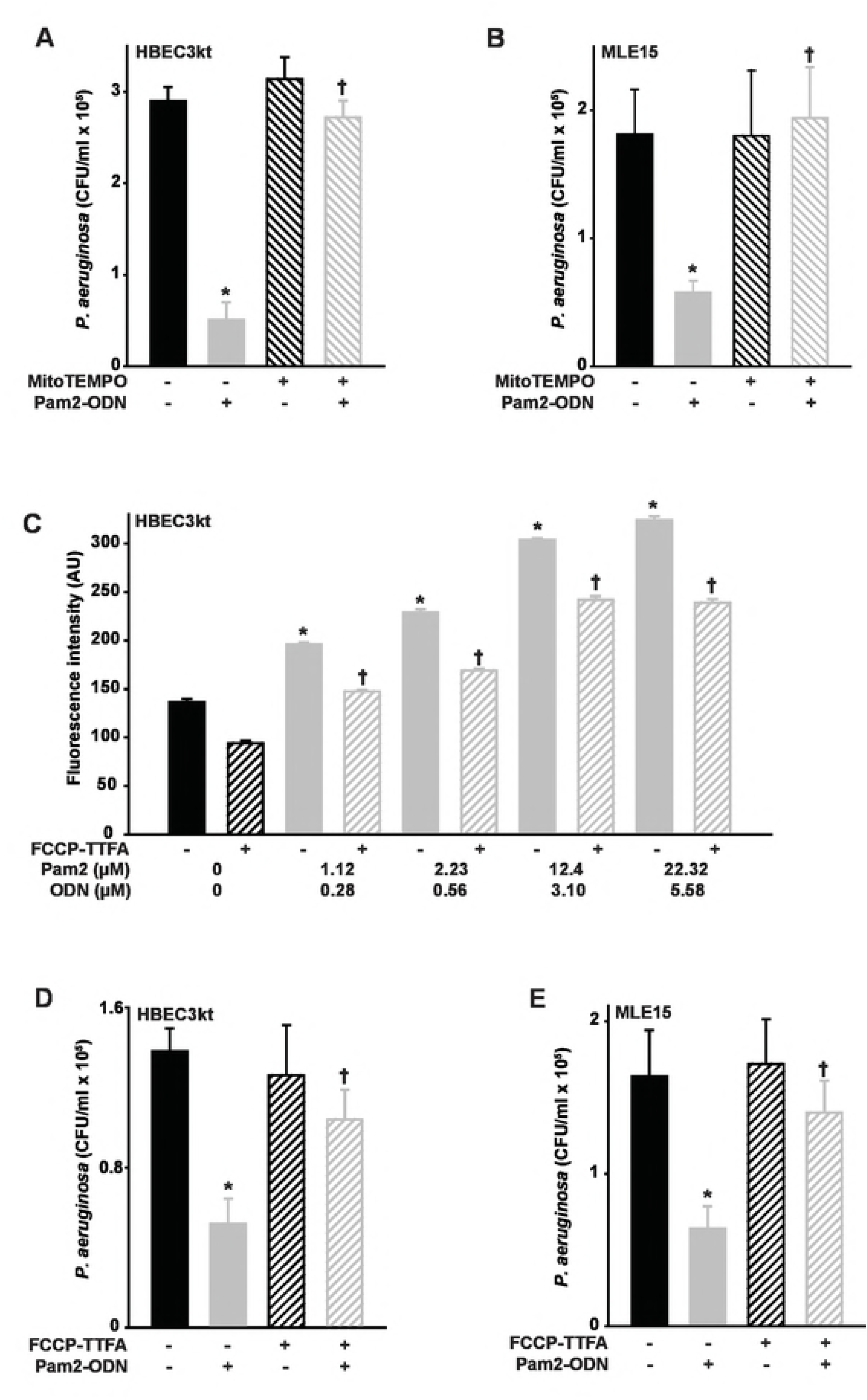
Mitochondrial ROS are required for Pam2-ODN-induced antibacterial epithelial responses. HBEC3kt (**A**) or MLE15 (**B**) cells were pretreated with MitoTEMPO or PBS, treated for 6 h with PBS or Pam2-ODN (Pam2 2.23 uM, ODN 0.56 uM), then challenged with *P. aeruginosa*. Shown are culture bacterial burdens 6 h after challenge. (**C**) HBEC3kt cells were pretreated with FCCP-TTFA or PBS, exposed to MitoSOX, then treated with PBS or Pam2-ODN at the indicated doses. Shown are culture fluorescence intensities at 100 min after treatment. HBEC3kt (**D**) or MLE15 (**E**) cells were pretreated with FCCP-TTFA or PBS, treated for 6 h with PBS or Pam2-ODN (Pam2 2.23 uM, ODN 0.56 uM), then challenged with *P. aeruginosa*. Shown are culture bacterial burdens 6 h after challenge. Each panels is representative of at least three independent experiments. N= 4-5 samples/condition for all experiments. * p < 0.01 vs. PBS-treated without inhibitor/scavenger; † p < 0.02 vs. Pam2-ODN-treated without inhibitor/scavenger.

## DISCUSSION

Although the airway and alveolar epithelia have historically been considered inert barriers, accumulating evidence now clearly supports their essential contributions to antimicrobial defense. In addition to their established capacity to recruit and activate leukocyte-mediated defenses in the lower respiratory tract, epithelial cells can exert directly antimicrobial effects on invading pathogens (22). Indeed, we have found that lung epithelial cells function as primary effector cells of inducible resistance to pneumonia, and we have reported the epithelial expression of numerous antimicrobial molecules following in vitro or in vivo exposure to Pam2-ODN (16, 17, 21, 23). The current work demonstrates that a single epithelial Pam2-ODN treatment promotes mouse survival of bacterial challenges by reducing pathogen burden and attenuating associated immunopathology. Similar in vivo Pam2-ODN-induced reductions in pathogen burden have been demonstrated by every investigated quantification technique, suggesting that the induced antimicrobial environment results in elimination of bacteria from all anatomic and cellular compartments of the lungs. This same inducible pathogen killing effect is observed from Pam2-ODN-treated isolated lung epithelial cells, where the reduced pathogen burden enhances cellular survival of bacterial challenges, even in the absence of leukocyte contributions.

Although numerous transcriptionally-regulated antimicrobial peptide species are induced by Pam2-ODN treatment of epithelial cells, no individual peptides have been demonstrated to be required for the Pam2-ODN-enhanced mouse survival of infectious challenge by any pathogen nor for the inducible intrapulmonary pathogen killing that uniformly correlates with inducible resistance. While it was, perhaps, unanticipated that none of these highly enriched peptides would prove essential to the protection, there are a number of plausible explanations for this observation. Foremost among these is the possibility that the extreme multiplicity of induced antimicrobial molecules renders the loss of only one or two peptides largely irrelevant, since the remaining species are sufficient to protect. Another important alternate consideration is that different peptide combinations may be required to protect against different challenges. So, we might eventually recategorize certain peptides that we have previously deemed to be non-essential for protection as required for protection against specific (as yet untested) alternate pathogens.

Regardless of explanation, the inability to detect any essential antimicrobial peptides, however, serves to underscore the robustness of the Pam2-ODN-induced protection against otherwise lethal pneumonias and profoundly emphasize the importance of our recent finding that epithelial ROS production is required for inducible antiviral defense. Unlike the peptide studies, we have clearly established that inducible ROS are essential to protecting against in vivo and in vitro challenges by orthomyxoviruses and paramyxoviruses (23). The present study similarly finds that inducible ROS production is essential to Pam2-ODN-induced epithelial bacterial killing. Strikingly, we find that ROS derived from both DUOX-dependent and mitochondrial sources are required for protection.

ROS have been long recognized to contribute to antibacterial defenses, particularly in the context of NADPH oxidase-dependent killing of bacteria in phagolysosomes of myeloid cells (26). However, the broadly microbicidal capacity of ROS generated by a wide range of cells has been demonstrated in recent years with evidence that ROS can exert antimicrobial actions against Gram-negative, Gram-positive, viral and fungal pathogens, even in the setting of established biofilms or antibiotic resistance (38-40). This vigorous protection may rely in part on the capacity of ROS-dependent strategies to synergize with other antimicrobial treatments to overcome antibiotic resistance (38, 41). Similarly, ROS can also promote bacterial clearance by reducing the minimum inhibitory concentrations of host effector molecules, such as neutrophil proteases (26). Consequently, it has been proposed that therapeutic induction of microbial ROS may potentiate the microbicidal activity of other antibacterials (42) and that eliciting ROS production is an essential component of the pathogen killing mechanisms of some antimicrobial pharmaceutical agents (43, 44).

Superoxide and hydrogen peroxide are the predominant species produced by lung epithelial cells (26, 34, 45). We have recently presented a comprehensive assessment of ROS production following Pam2-ODN treatment of isolated lung epithelial cells. Congruent with the findings in the current work, we found superoxide and hydrogen peroxide to be the only species detectably induced by Pam2-ODN (23).

The principal sources of hydrogen peroxide from the lung epithelium are DUOX1 and DUOX2 (30, 31, 46, 47), and therapeutic induction of DUOX2 has been proposed as a potential antimicrobial therapeutic strategy (48). Indeed, we have reported that DUOX-related genes are enriched following epithelial exposure to Pam2-ODN, both in vitro and in vivo (17, 23). The current study confirms that DUOX-dependent ROS are required for inducible *P. aeruginosa* killing, as this protective effect is profoundly attenuated when cellular ROS are annihilated by PEG-HCC or either DUOX is selectively knocked down. It is well established that the DUOX-dependent product of the lactoperoxidase/hydrogen peroxide/thiocyanate system, hypothiocyanate, exerts antimicrobial effects (46, 47, 49), however it is unlikely that the ROS dependency of Pam2-ODN-induced protection simply reflects hydrogen peroxide-mediated hypothiocyanate production, as our in vivo models lack tracheobronchial seromucus glands as a lactoperoxidase source (50) and our in vitro models lack a source of thiocyanate (33). This suggested that the DUOX-dependent ROS effects are likely achieved either through direct pathogen toxicity or through host signaling events.

In addition to DUOX-dependent hydrogen peroxide production, mtROS have been increasingly reported to contribute to wide ranging aspects of both innate and adaptive immunity (34, 51), and increased production of mitochondria-derived species likely explains the Pam2-ODN-increased superoxide we have detected here and in previous work (23).

mtROS are generated via leakage from the electron transport chain (45), resulting in production of superoxide that diffuses through mitochondrial membranes following dismutation to hydrogen peroxide (52). This process is exquisitely tightly regulated by changes in scavenging, production, and localization (34), so the substantial induction of mtROS by Pam2-ODN represents an important homeostatic perturbation. A reported eleven different mitochondrial sites can be perturbed to result in mtROS production and release (53), and we are presently investigating the signaling events that promote Pam2-ODN-induced mtROS production. However, it has been reported that TLR manipulation can promote generation of both antibacterial mtROS (37) and NADPH oxidase-generated ROS (54) in macrophages, so it is highly plausible that TLR ligands could induce ROS from both mitochondrial and DUOX sources in epithelial cells. Interestingly, although coordinated regulation of NADPH oxidase ROS and mtROS production has been reported (52, 55), the current study and our recent antiviral work (23) remain the only reports of concurrent induction of ROS from mitochondrial and DUOX sources in any cell type.

The precise mechanisms underlying the unanticipated requirement for dual sources of ROS have yet to be elucidated. However, this may be explained by dependence on the phenomenon of ROS-induced ROS to promote high ROS concentrations (52, 56, 57). Alternate potential explanations also include the hypothesis that multiple sources are required to achieve sufficiently high aggregate ROS concentrations to exert microbicidal actions (additive effect) or that the different sources play different roles, such as one source directly causing pathogen membrane damage while the other facilitates ROS-dependent signaling events. It is even possible that the mitochondrial function serves to regulate DUOX functions (58). This remains an area of active investigation.

These data indicate that multisource ROS are required for Pam2-ODN-induced bacterial killing, extending the range of pathogens that are known to be susceptible to inducible epithelial ROS and highlighting the centrality of ROS generation to the protective phenomenon of inducible epithelial resistance. By advancing understanding of the mechanisms of Pam2-ODN-induced resistance, these data may facilitate development of even more efficacious resistance-inducing therapeutics and offer hope that pneumonia can be prevented in vulnerable populations.

## MATERIALS AND METHODS

### Animals, cells, and reagents

All general reagents were obtained from Sigma-Aldrich (St Louis, MO), except as indicated. All mouse experiments were performed with 5–8 week-old C57BL/6J (The Jackson Laboratory, Bar Harbor, ME) of a single gender in accordance with the Institutional Animal Care and Use Committee of The University of Texas MD Anderson Cancer Center, protocol 00000907-RN01. Immortalized human bronchial epithelial (HBEC3kt) cells were kindly provided by Dr. John Minna. Murine lung epithelial (MLE-15) cells were kindly provided by Dr. Jeffrey Whitsett. The cell lines used were authenticated by the MD Anderson Characterized Cell Line Core Facility.

### Cell Culture

HBEC3kt cells were cultured in keratinocyte serum-free media (KSFM, ThermoFisher Scientific, Grand Island, NY) supplemented with human epidermal growth factor and bovine pituitary extract. MLE-15 cells were cultured in RPMI supplemented with 10% fetal bovine serum. Cultures were maintained in the presence of penicillin and streptomycin.

### TLR treatments

For *in vivo* studies, S-[2,3-bis(palmitoyloxy)-propyl]-(R)-cysteinyl-(lysyl) 3-lysine (Pam2CSK_4_) and ODN M362 (InvivoGen, San Diego, CA) were reconstituted in endotoxin-free water, then diluted to the desired concentration in sterile PBS. As previously described (16), the Pam2-ODN was placed in an Aerotech II nebulizer (Biodex Medical Systems, Shirley, NY) driven by 10 l min^-1^ air supplemented with 5% CO2 for 20 min. The nebulizer was connected by polyethylene tubing to a polyethylene exposure chamber. 24 h prior to infections, 8 ml of Pam2 (4μM)-ODN (1μM) was delivered via nebulization to unrestrained mice for 20 minutes, and then mice were returned to normal housing. For *in vitro* studies, Pam2-ODN was added to the culture media 4 h prior to inoculation with bacteria or at the indicated time point. Pam2-ODN was given in fixed ratio, but at varying doses as indicated.

### Infection models

As previously described (16), frozen stock of *Pseudomonas aeruginosa* strain PA103 (American Type Culture Collection, Manassas, VA) was incubated overnight in tryptic soy broth, then expanded in Luria-Bertini media to OD_600_ 0.35. Bacterial suspensions were centrifuged, washed, re-suspended in PBS, and aerosolized over 60 min. For all bacterial challenges, a nebulized inoculum of 10 ml of ~2 x 10^10^ CFU/ml were delivered. Immediately after bacterial challenges, some mice were anesthetized and their lungs were harvested and homogenized (16) using a Mini-Beadbeater-1 (Biospec, Bartlesville, OK). Serial dilutions of the nebulizer inoculum and lung homogenates were plated on tryptic soy agar plates (Becton Dickinson). The remaining mice were observed for 12 d to determine whether their clinical conditions met euthanasia criteria. Following infection, lab personnel coordinated with staff of the MD Anderson Department of Veterinary Medicine to ensure that the mice were evaluated a minimum of three times daily to determine whether euthanasia criteria were met. As specified in the above noted animal protocol, mice that were not submitted to anesthetic excess followed by thoracotomy with bilateral pneumonectomy for pathogen burden assessments were humanely sacrificed by inhalational exposure to approved concentrations of carbon dioxide until respiratory efforts ceased, followed by cervical dislocation as a secondary method of euthanasia, when they either achieved the end of the observation period or met the predesignated euthanasia criteria. The relevant euthanasia-triggering criteria include any evidence hypothermia, impaired mobility, respiratory distress, inability to access food or water, or any evidence of distressed behavior. Weight loss is also among the approved indications for euthanasia, but the mice that met euthanasia criteria in this model became ill or distressed within 1-2 d (before losing > 25% body weight), so no mice were euthanized due to weight loss in the current study. Despite the close observation, this same rapidity of illness resulted in up to 3 of the 56 infected mice dying spontaneously from pneumonia before being euthanized in some experimental replicates. Although meeting euthanasia criteria is the primary endpoint, the presented “Survival (%)” in Figure 1 formally indicates mice that had not either met euthanasia criteria or spontaneously died. When mice were identified to meet criteria, they were submitted to euthanasia within 30 minutes by either lab personnel or Department of Veterinary Medicine staff. All lab personnel and Department of Veterinary Medicine staff receive formal instruction in methods to minimize stress and discomfort to experimental animals and analgesia is provided to animals that demonstrate any evidence of discomfort but do not meet euthanasia criteria.

For the in vitro challenges, after the indicated treatments, confluent mouse or human epithelial cell cultures were inoculated with *P. aeruginosa* (20 μl − 1×10^5^ CFUs/ml), incubated for 6 h, then harvested and submitted to serial dilution culture.

### *Lentiviral shRNA knockdown of* DUOX1 *and* DUOX2

GIPZ human *DUOX1* and *DUOX2* lentiviral shRNA clones were purchased from GE Dharmarcon (Lafayette, CO). Lentiviruses bearing human *DUOX1* and *DUOX2* shRNA were produced by transfection in 293T cells per manufacturer’s instructions. Infection efficiency was enhanced by addition of 8 μg/ml Polybrene into the culture media and centrifuging the cells at 2,000 rpm for 60 min at 32 °C. Lentiviral-infected HBEC3kt cells were selected by cell sorting based on GFP expression. shRNA knockdown efficiency was determined by immunoblot analysis, as previously shown (23).

### ROS detection, scavenging and inhibition

To assess ROS generation, cells were treated with 5μM of each indicated detector for 1 h before exposure to Pam2-ODN or sham, as previously reported (23). Fluorescence was continuously measured on a BioTek Synergy2 for 250 min after treatment. Excitation/emission wavelengths for ROS-detecting agents are: Carboxy-2’,7’-dichlorodihydrofluorescein diacetate (CO-H_2_DCFDA, ThermoFisher), 490nm/525nm; and MitoSOX^™^ Red (ThermoFisher), 510nm/580nm.

Cellular ROS were scavenged by 1 h exposure to PEGylated hydrophilic carbon clusters (PEG-HCC, 5μg/mL) prior to application of Pam2-ODN or PBS (23). Mitochondrial ROS were scavenged by 1 h exposure to (2-(2,2,6,6-Tetramethylpiperidin-1-oxyl-4-ylamino)-2-oxoethyl) triphenylphosphonium chloride monohydrate (MitoTEMPO, 30nM, ThermoFisher) prior to treatment with Pam2-ODN or PBS (23). Disruption of *in vitro* mitochondrial ROS production was achieved through concurrent application of trifluoromethoxy carbonylcyanide phenylhydrazone (FCCP, 400 nM, Cayman Chemical, Ann Arbor, MI), and 2-thenoyltrifluoroacetone (TTFA, 200 μM, Sigma) (23).

### Statistical analysis

Statistical analysis was performed using SPSS v19 (SAS Institute, Cary, NC). Student’s t-test was used to compare the lung pathogen burdens between the groups. Error bars shown in all the figures represent technical replicates within the displayed experiment, rather than aggregation of experimental replicates. Percentage of mice surviving pathogen challenges was compared using Fisher’s exact test on the final day of observation, and the log-rank test was used to compare the survival distribution estimated by the Kaplan–Meier method.

## ACKNOWLEDGEMENTS

This study was supported by NIH grants R01 HL117976 and DP2 HL123229 to S.E.E., and P30 CA016672 to the MD Anderson Cancer Center.

## SUPPORTING INFORMATION FIGURE LEGENDS

**Figure S1. Early and late bacterial burden reduction by Pam2-ODN treatment**. Wild type C57BL/6J mice were treated with Pam2-ODN or PBS (sham) 24 h prior to challenge with fluorescently-labeled P. aeruginosa. (**A**) Representative micrograph overlays of lung sections immediately after infection and 24 h after infection. Blue = DAPI, Green = bacteria. (**B**) Quantification of GFP signal in the indicated conditions. N = 3 mice per condition, 10 high power fields measured per mouse. * p = 0.002 vs. PBS treated 0 h after challenge. p = 0.00006 vs. PBS treated 24 h after challenge.

**Figure S2. Cytokine and antimicrobial peptide induction in Pam2-ODN-induced resistance**. (**A**) HBEC3kt cells were treated with PBS (sham) or Pam2-ODN for 2 h, then submitted to RT-qPCR for the indicated transcripts. Shown are RQ values for the target transcript relative to 18s gene. Each panel is representative of at least three independent experiments. N=4-5 samples/condition for all experiments. Wild type or mice deficient in (**B**) *Lcn2*, (**C**) *Cramp*, (**D**) *Lcn2* and *Cramp*, or (**E**) the indicated acute phase SAA genes were treated with PBS (sham) or Pam2-ODN by aerosol 24 h prior to challenge with *P. aeruginosa*. Shown are survival plots for each challenge. Each panel is representative of at least three independent experiments. N=8-10 mice/condition. * p < 0.001 vs PBS treated. ** p < 0.007 vs. syngeneic PBS treated. † p < 0.05 vs. syngeneic PBS treated.

